# Calibrated structural homology transfer yields putative molecular functions for domains of unknown function in four model proteomes

**DOI:** 10.64898/2026.09.11.750891

**Authors:** Jodh Singh Pannu, Kendall Green, Jeffrey Vedanayagam

## Abstract

Improvements in computational protein structure prediction have enabled searches for remote homologs of proteins whose molecular function remain unknown. However, the reliability of functional annotations from such searches has not been systematically quantified. In this study, we assembled a time-split benchmark from Pfam families annotated as domains of unknown function (DUFs) in Pfam 28.0 and were subsequently assigned a function in the current release (Pfam 38.0). These retrospective-DUFs provide ground truth for assessing functional annotation from remote homology searches. In our benchmark analysis, we paired retrospective-DUFs with a difficulty-matched arm from known domains to distinguish query difficulty from the method’s performance. Across four model proteomes (yeast, *C. elegans*, *Drosophila*, and mice), Foldseek searches against AlphaFold/Swiss-Prot, PDB100, and CATH50 recovered the later-assigned function for 15.9% of retrospective DUF queries, compared with 30.1% of matched known-domain queries, after masking for self-family and self-clan level hits to remove circularity. At a fixed confidence cut-off (qTM ≥ 0.5), 55.4% of informative calls on retrospective DUFs were incorrect, establishing an error model for prospective use. Furthermore, in our comparison of tools for homology searches, Foldseek outperformed MMseq2-based sequence search but was not statistically separable from the ESM-2 protein language model embedding baseline. Applying the calibrated structural homology search pipeline to 296 currently unannotated DUF queries in Pfam 38.0 yielded 50 confident, specific functional assignments. We provide the benchmark and ranked candidates as a resource to facilitate functional studies.

## Introduction

A substantial fraction of protein-coding genes in prokaryotic and eukaryotic proteomes lack assigned molecular functions (1). Across four extensively studied model eukaryotes (*Saccharomyces cerevisiae*, *Caenorhabditis elegans*, *Drosophila melanogaster*, and *Mus musculus*), 29.7% of annotated proteins lack molecular function and curated functional description ranging from 12.3% in *S. cerevisiae* to 51.1% in *C. elegans* (2, 3). Since unveiling the genomes of major model organisms nearly two decades ago (4–7), the fraction of genes with unknown molecular function has remained more or less stable rather than shrinking. Most research has focused on already characterized genes, so little attention has gone to the uncharacterized remainder (8). Importantly, when studies have subjected proteins of unknown molecular function to systematic loss-of-function in mutant screens, many have proved to be indispensable. For instance, mutant screens across thousands of bacterial genes of unknown function have recovered important phenotypes (9), and domains of unknown function (DUFs) were found to be essential in bacteria at rates comparable to those of well-characterized protein domains (10). Recent efforts, dubbed ‘functional unknowmics’, have also systematically screened conserved proteins of unknown molecular function (11) and uncovered crucial phenotypes, emphasizing that the unknown proteome is an important yet-to-be-resolved reservoir of biology.

Systematic sequence-based classification of protein domains led to the development of the Protein Families (Pfam) Database (12), and Pfam’s ‘domain of unknown function’ (DUF) nomenclature makes the gap in our understanding the molecular functions of these domains explicit. DUFs share a recognizable and evolutionarily conserved domain for which no function has been determined (12–14), and the defined domain boundaries slightly vary among databases such as Pfam (13) and SMART (15). Nevertheless, the availability of family boundaries for these DUFs makes them candidates for attempting computational functional assignment (16). The traditional approach to remote homology detection was sequence-based. For instance, PSI-BLAST (17), profile-profile comparison (18), and hidden Markov model procedures (19) allowed assignment of DUF to a characterized protein family when sequence similarity is too weak for pairwise searches (20, 21). However, these methods lose discriminating power below 20-25% sequence identity, which is precisely the realm DUFs occupy.

Beyond sequence-level searches, protein 3D structure can, in principle, reach further in detecting remote homology. Protein fold similarity is known to persist even when sequence-level similarity has decayed, and structure-based homology detection has a long history of detecting remote homology that sequence-based methods fail to recover (22–24). However, what has changed in recent years is the coverage of search databases and the speed at which structural alignments can be performed. AlphaFold2 generated near-accurate 3D structure models available at proteome scale (25, 26), and the AlphaFold Protein Structure Database (AFDB) now comprises > 200 million predicted structures (27). On the search tools end, Foldseek made structural similarity searches fast and efficient (28), and clustering the predicted structural universe has already uncovered novel families and folds with no prior structural representatives (29, 30). These efforts have also made domain-level classification of whole predicted proteomes possible (31, 32). In addition to structure-driven alignment modes, recent developments in protein language models offer an additional arm to identify domain similarity by transferring annotations from embedding space without an explicit alignment (33, 34). Together, these developments make a structure-first approach to tackle uncovering functional roles for DUFs technically straightforward and feasible.

While structural similarity searches can be efficient with new tools, executing them and trusting them are two different problems, and the second one remains unresolved. A structural hit reveals shared ancestry or shared fold. However, it is not evidence of shared molecular function, and annotation transfer is a step where errors can occur. It is also known that function does not track protein fold cleanly. For instance, homologous enzyme superfamilies diverge in catalytic activity, and annotation transfer between them is a well-known source of systematic mis-annotation in public databases (35). Critically, once introduced, such errors easily propagate through databases by the same annotation transfer mechanism that created them (36, 37). Indeed, community assessments of function prediction have attempted to quantify how hard the general problem is (38, 39), but those evaluations measure prediction for proteins that already have annotations to withhold, which is the opposite for the DUF case, where there is little to no ground truth available to evaluate for DUF proteins. As a result, it is difficult to quantify how often the transferred function is correct for DUFs.

Furthermore, two additional factors compound the problem. First, predicted models carry confidence that varies within a chain, and low-confidence or disordered regions support neither good superposition nor reliable interpretation (26, 40). Second, any benchmark built by searching a database that contains the query’s own family measures retrieval and not function transfer. This circularity could inflate apparent performance without being visible in the output. Therefore, to minimize circularity, our study reports a calibrated error rate for structure-based annotation transfer, specifically for DUFs. Here, we performed that calibration on retrospective DUFs (families that were labeled DUFs in Pfam 28.0 and have since been assigned functions in Pfam 30.0), and then apply it to still DUFs (DUFs that have no assigned functions in Pfam 30.0). To achieve that, we constructed a time-split benchmark with ground truth and use it to measure how often Foldseek-derived annotation transfer recovers the function a family was later given, using 353 retrospective DUFs in 4 model query organisms.

We paired retrospective DUF searches with a difficulty-matched arm of known domain controls, and the hits carrying the query’s own Pfam family or clan sibling were masked, symmetrically in both arms. Without masking for circularity, top-1 agreement is 84.7%; however, with it, the agreement falls to 26.1% for retrospective DUFs. We report error rates stratified by structural similarity and also by the specificity of the transferred molecular function term, using information content to discriminate a confident claim from an informative one (41). Finally, we applied the calibrated pipeline to the 296 DUF families that remain unannotated, yielding 50 confident, specific functional hypotheses with associated expected error rates for experimental follow-up studies.

## Results

### A retrospective benchmark pipeline and properties of DUF proteins in four model proteomes

In four focal proteomes in this study (*S. cerevisiae*, *C. elegans*, *D. melanogaster*, and *M. musculus*), nearly 6-10% of proteins with unknown molecular function contain a DUF/UPF (unknown protein function) family are listed in Pfam (13) (**Fig 1A**), which are a subset of ∼20% of protein-coding genes in these species with no known molecular function (**Fig S1A**). In addition to DUF families, several proteins with unknown molecular functions lack Pfam domain annotations. Whereas 24-65% of proteins have a domain content recognizable and labeled as a “named Pfam domain”, but their molecular function remains unassigned (**Fig. 1A**). The key motivation for this study is to use AlphaFold-predicted structures of DUF/UPF proteins to conduct structural homology searches and identify putative functional roles for yet-unresolved DUF families in Pfam. To develop a pipeline and conduct structural homology searches for DUF proteins, we first evaluated whether the predicted structures of DUF proteins are of comparable quality of proteins with a known molecular function. We assessed the overall quality of the predicted structures with AlphaFold’s predicted Local Distance Difference Test (pLDDT) (25). On average, the predicted structures for proteins of unknown molecular function have lower pLDDT scores, with ∼50% of predicted structures below pLDDT = 70 (high quality), compared to 20-25% for proteins with known function in the model proteomes (**Fig. 1B**).

**Figure 1:**
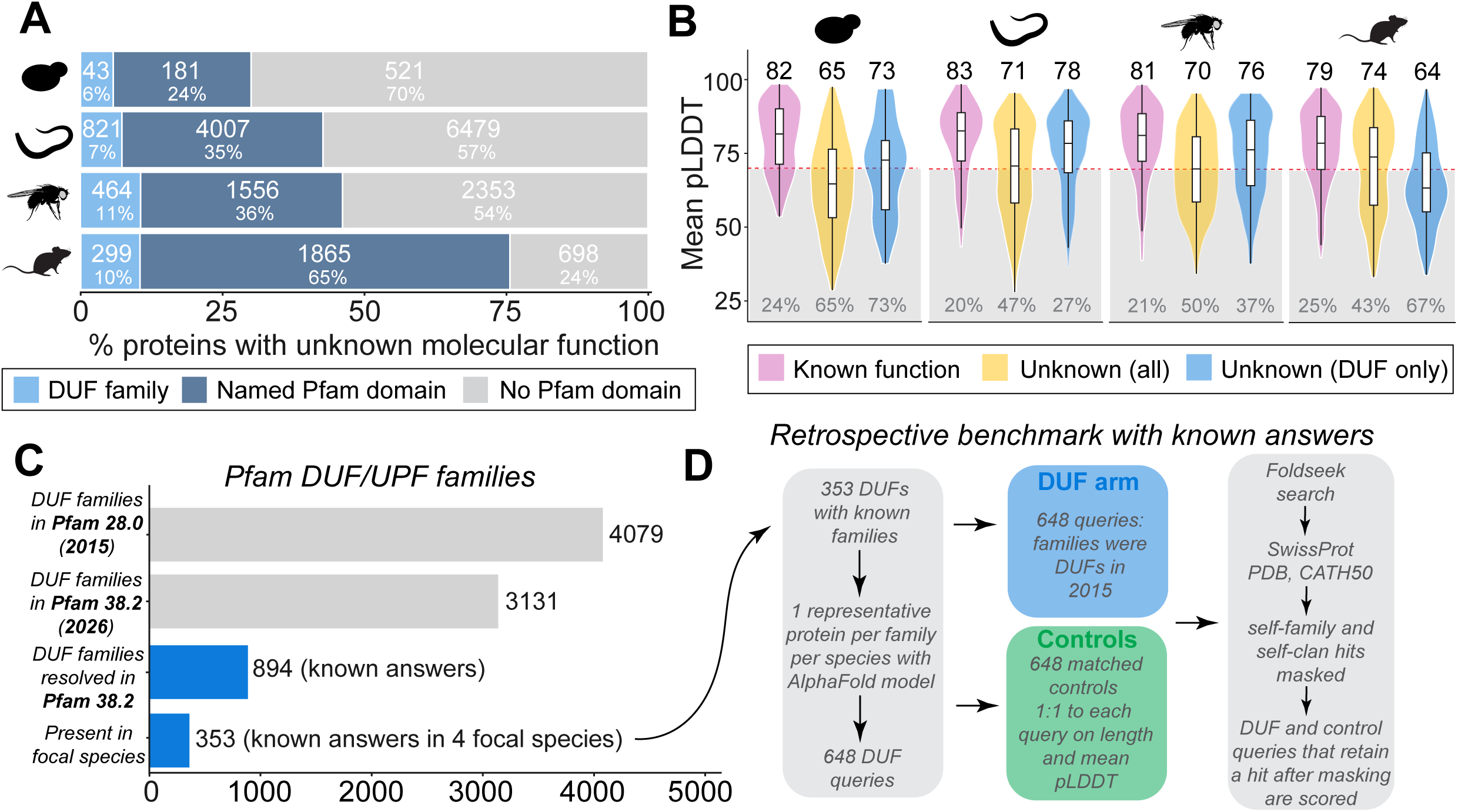
Benchmark pipeline and properties of proteins of unknown function. A) Proteins of unknown molecular function make up significant fraction of four focal model proteomes in this study. Specifically, proteins annotated to contain a DUF range from 6% in yeast to 11% in *Drosophila* proteome. B) Structural properties as measured by AlphaFold’s pLDDT score. Proteins separated as ones with known functional domain and unknown function show higher median pLDDT > 70 (high-confidence) for proteins with a known functional domain in all 4 model proteomes. A side-by-side comparison of pLDDT distribution of all proteins of unknown function, as well as proteins with a DUF are shown. Overall, the proteins of unknown function have median pLDDT values much lower than proteins with known function. When proteins with a DUF are separately considered, their median pLDDT scores are above 70 (high-confidence) in three or the four model proteomes, indicating good quality AlphaFold predictions for proteins with a DUF. C) Comparison of DUF families in Pfam between 28.0 and 38.0 releases. 2026 update has 894 DUF families resolved, of which 353 occur in the 4 model proteomes considered in this study for a time-split benchmark construction. D) Pipeline for retrospective benchmark construction for 353 DUF families, along with a difficulty-matched control arm with proteins containing a known functional domain. Structural homology searches were conducted using Foldseek, and a self-family and self-clan masking was implemented to minimize circularity after retrieval for functional annotation transfer.

Specifically for proteins containing DUFs, the average pLDDT is higher, with good, predicted structures in *C. elegans* and *D. melanogaster* (27-37% below pLDDT < 70), whereas 67-75% of DUF proteins have a predicted average pLDDT < 70 in *S. cerevisiae* and *M. musculus* (**Fig. 1B**). Furthermore, DUF proteins are typically half the length of proteins with a known function, where the median length of DUF proteins in four model proteomes range from 133-250 amino acids (aa), while the length range for proteins with a known domain/function is 401-456 aa (**Fig. S1B**). We conducted these analyses to identify difficulty-matched, length-controlled known domain arms for our benchmark efforts. From the four model proteomes, we first identified DUF families that were resolved for a molecular function between the Pfam 28.0 and 38.2 releases to construct a time-split benchmark. Of the 894 DUF families that were resolved in the new Pfam release, 353 DUFs reside in the four focal species (**Fig. 1C**), providing ground truth to assess for the structural homology searches. From these, we identified one representative protein per DUF family per species, yielding 648 DUF queries for the benchmark pipeline (**Fig. 1D**; **Supplementary Table S1**). Alongside the DUF query arm, we also identified 648 length- and mean pLDDT-matched controls with a known functional domain to evaluate the benchmark pipeline and compare query difficulty (DUF vs. known domain) and outcomes from structural homology searches. We utilize Foldseek (28) for structural homology searches and provide a comparison of Foldseek’s performance to sequence-based homology searches such as MMseq2 (20) as well as the ESM-2 protein language model (34) in our evaluation of the pipeline (described in the following section). Briefly, our pipeline (**Fig. 1D**) searched the retrospective DUFs and matched known domain controls against three databases, and the resulting output was masked for circularity (see Methods). The hits that survived after masking were then scored using the InterPro-GO function. A detailed subsection of the pipeline that involves masking and scoring procedures are provided in **Fig S2A and B**.

### Benchmarking on retrospective DUFs provide yield rates for prospective use on domains of unknown function

We used three different databases (AFDB/SwissProt, PDB100, and CATH50) for Foldseek searches (**Fig. 2**). Each query in both the retrospective DUF and known domain arms was searched against these databases, which represent a wide range of database sizes and annotation densities. AFDB/SwissProt is well-curated and represents >540,000 proteins that are near-completely annotated (42). PDB100 has a database size of >340,000 proteins, including experimentally resolved protein structures (43), while CATH50 is a structural classification database that assigns domains to the structures in PDB (5841 domain superfamilies) (44). Throughout the search pipeline, the resulting hits from all three databases for query proteins were self-family and self-clan masked, and a ‘call’ described below is the functional term transferred from the top-ranking surviving hit, and we assert a call is scored correct if it agrees with Pfam family assignment for both retrospective DUFs and known domain controls.

**Figure 2:**
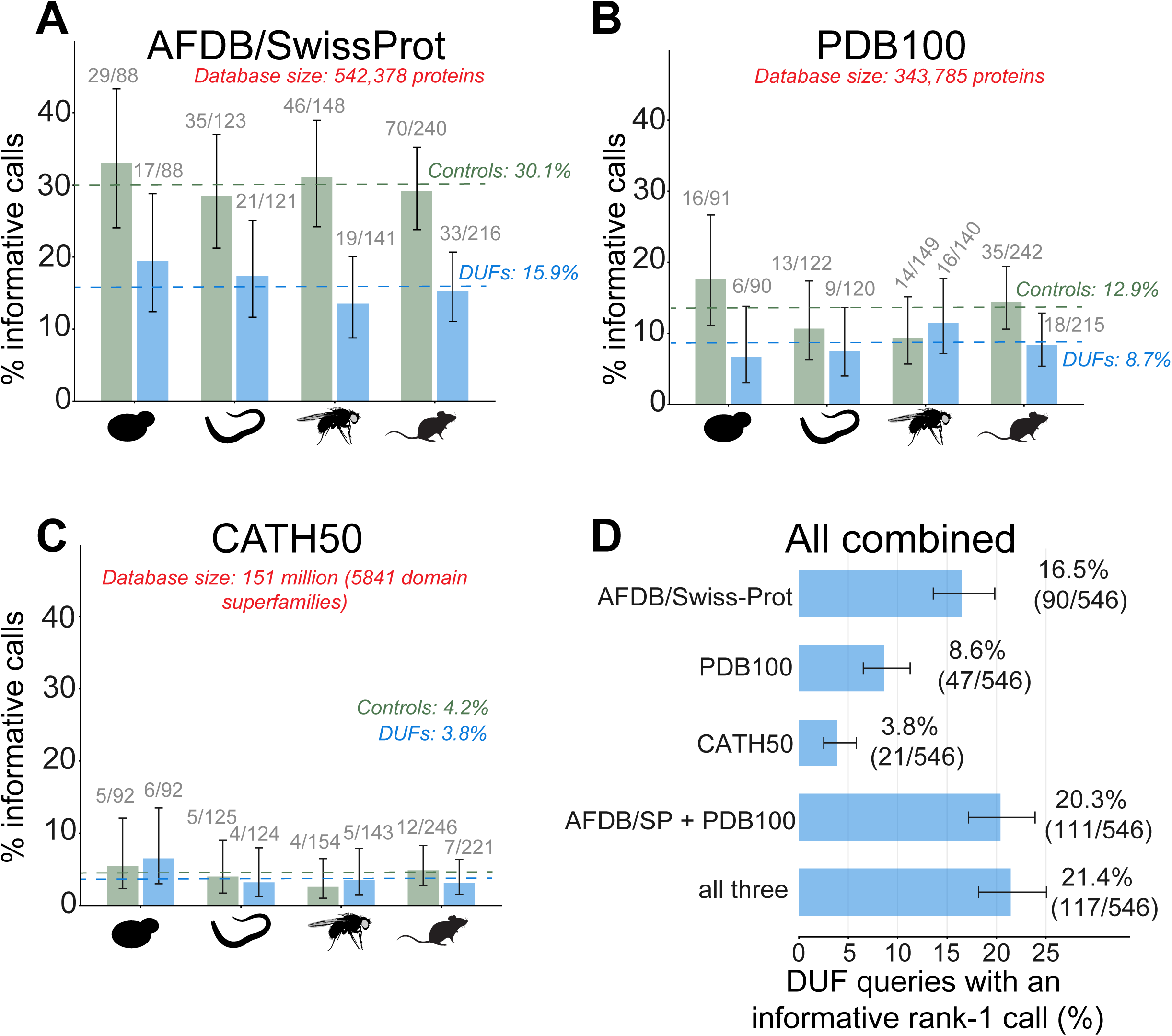
Time-split benchmark comparison between retrospective DUFs and known domain controls. 648 queries from both retrospective DUFs and known domain control arms were searched against three databases (AFDB/SwissProt, PDB100, and CATH50) using Foldseek. A) AFDB/SwissProt fetched most informative calls out of the three databases and 15.9% of the retrospective DUF top-1 calls were correct, compared to 30.1% for the control arm in 4 model proteomes. B and C) Similar searches against PDB100 and CATH50 recovered 8.7% and 3.8% of correct top-1 rank calls for retrospective DUFs, respectively. D) All three databases combined produced 21.4% (117/546) correct top-1 rank calls and the addition was complementary and non-overlapping entries were contributed by PDB100 and CATH50. Rank-1 calls are the best e-value hit after masking at self-family and self-clan levels, and error bars are 95% Wilson score intervals for a binomial proportion (z = 1.96, no continuity correction, unadjusted for multiple comparisons).

For our searches against AFDB/SwissProt, the known domains control arm received a correct rank-1 call for 30.1% (180/599 queries from four species), whereas the retrospective DUFs arm identified a correct rank-1 call for roughly half the rate: 15.9% (90/566) queries (**Fig. 2A**). In contrast, the PDB100 and CATH50 returned a correct rank-1 call for 12.9% and 4.2% for the control arm, and 8.7% and 3.8% for the retrospective DUFs arm, respectively (**Fig. 2B** and **2C**). For the retrospective DUF arm, the occurrence of a correct answer in the top-5 ranks increased the correct call to 23.5% for AFDB/SwissProt, 18.8% for PDB100, and 14.1% for CATH50 (**Fig. S3A**). The known-domains control arm outperformed the retrospective DUF arm across all databases, as expected for queries whose fold is better represented among annotated targets. In addition to these three databases, we also conducted a structural homology search against the AFDB/UniProt 50 (AFDB50) database, which contains > 52 million entries; however, AFDB50 yielded ∼30-fold fewer informative calls than AFDB/Swiss-Prot despite retrieving more hits per query, because circularity masking removes annotated family-carrying targets, leaving uninformative entries occupying top-ranked hits (**Fig. S3B**).

At the per-organism level, the known domain controls outperform retrospective DUFs across all databases, except for a change in direction in which DUFs score above controls in *D. melanogaster* against PDB100 (11.4% *vs*. 9.4%). However, the confidence intervals overlap, and we interpret this as sampling variation rather than a real reversal. Combining the informative calls from all three databases, we find that the results obtained from multiple databases are complementary, and overall, the correct call at top-1 rank for retrospective DUFs increases to 21.4% (117/546 queries) (**Fig. 2D**). Adding PDB100 and CATH50 rescued 27 DUFs that AFDB/SwissProt alone misses, 21 from adding PDB100 and 6 from adding CATH50. Only 26 of the 90 AFDB/SwissProt calls were also made by PDB100 and CATH50, confirming that the increment obtained from including multiple databases reflects a genuinely different retrieval than redundant confirmation (**Supplementary Table S2**). Taken together, our findings show that AFDB/SwissProt is the primary annotation transfer for informative calls, with PDB100 as a worthwhile secondary search, and CATH50 restricted to fold-level corroboration.

Finally, to confirm that the masking procedure drives the yield rates of rank-1 calls reported, we re-scored the same AFDB/SwissProt hits with masking disabled. Without masking at the self-family and self-clan levels, the top-1 functional agreement rises to 84.7% in the retrospective DUF arm and 84% in the known domains control arm. Masking at the Pfam family level alone reduces the agreement to 29.1% and 46.2% for retrospective DUFs and known domain controls, respectively, and adding clan-level masking further reduces this to 25.6% and 34.2% informative rank-1 calls (**Fig. S3C**). The unmasked rates are inflated by construction as 96% of unmasked top-1 hits carry the query’s own Pfam family or clan, so the unmasked pipeline is largely retrieving the answer it was asked to predict. The removal is also asymmetric between the known domain controls and retrospective DUF arms, as family-and-clan level masking discards 58.4% of control hits but 42.2% of DUF hits, because the difficulty-matched control families have more annotated close relatives (at the clan level) to retrieve. Therefore, the central comparison of this study exists only after the circular hits are removed.

### Evaluation of ground truth from retrospective DUFs and benchmark error rates

Despite an informative call, there are two likely failure modes with different consequences for interpreting the transfer of function. First, a call can be wrong because the transferred term is unrelated to the protein’s actual function, or it can be technically correct but so close to the root of the GO hierarchy that the information obtained is too general and not specific. We graded the hits that survived circularity masking into three tiers (**Fig. 3A**). First, homology searches obtain excellent structural alignment, but the target carries no molecular function annotation. For example, our confident-only hit for PF02696 shows a structural alignment in PDB100 (6iny; qTM 0.67; 27.3% identity) whose function is unknown. PF05907 illustrates the second outcome: its hit transfers zinc-ion binding (AF-O60140, qTM 0.63, 17% identity; confident + informative class) with an information content score of 4.0, which is correct but too general. Finally, PF06155 illustrates the case our approach is intended for, where the structure homology search fetches the top-ranked hit as γ-butyrobetaine dioxygenase (AF-Q19000, qTM 0.72, 16.6% sequence identity; confident + specific class), a specific enzymatic activity recovered at a sequence identity far below what conventional sequence-only based approaches typically use to identify homology.

**Figure 3:**
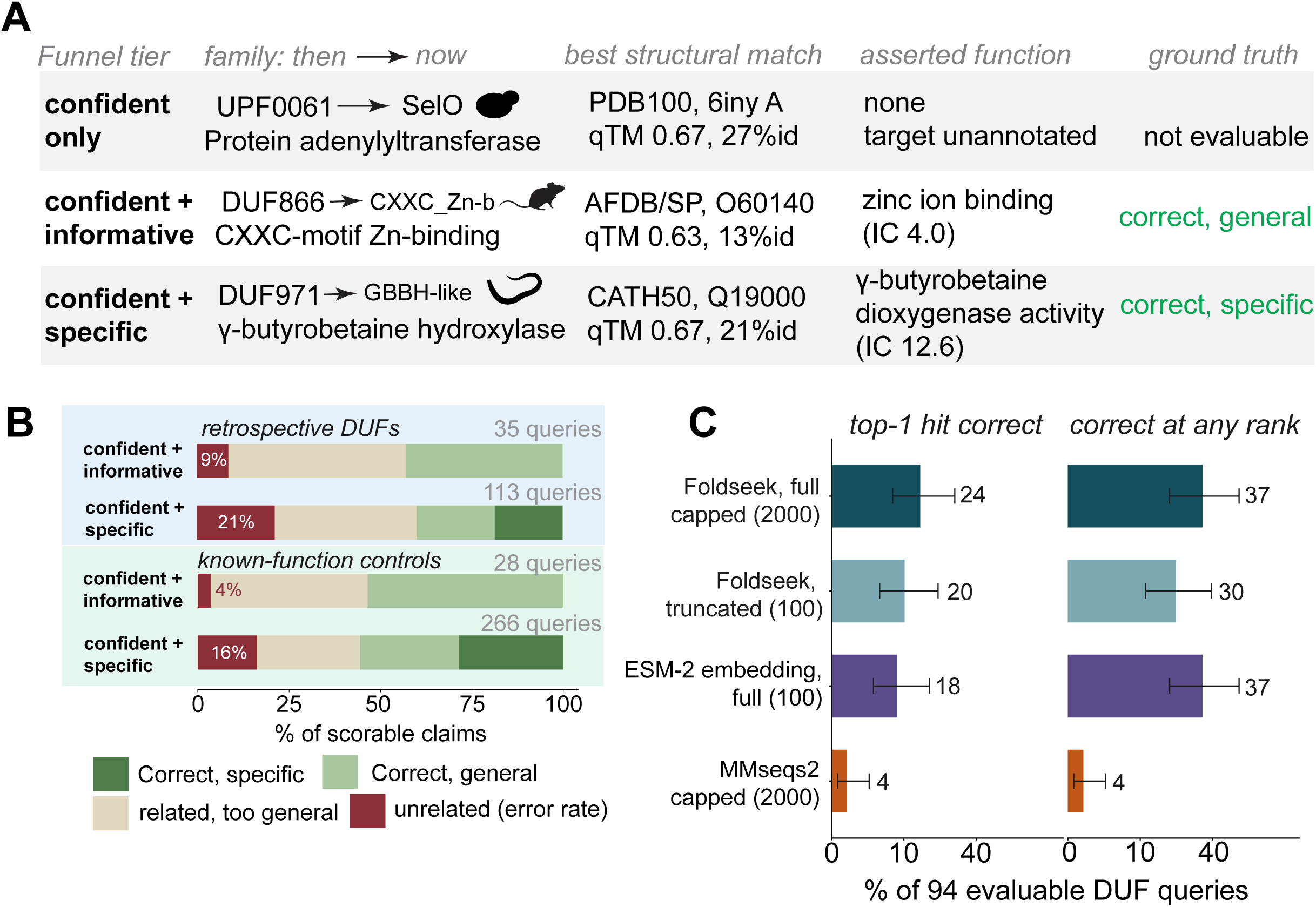
Time-split benchmark error rates and evaluation of Foldseek’s performance against embedding and sequence-based methods. A) Illustrative examples for three different funnel tiers for the hits based on how well top-1 hits conform to ground truth. Confident only tier has no transferable function despite good structural similarity and therefore no evaluable ground truth. The confident + informative tier has an asserted function, but the information content score is too general and the ground truth class in interpreted as correct, but general function. The third funnel tier is confident + specific transferred function, which high IC score identifying specific function as the ground truth. B) Distribution of confident + informative and confident + specific calls for the retrospective DUFs and the known domain control arms. Of the % scorable claims, 9-21% of the claims were unrelated hits compared to ground truth for the retrospective DUF arm, whereas 4-18% of the claims were unrelated for the known domain controls arm. C) Comparison of Foldseek’s performance against ESM-2 embedding based functional transfer and sequence-based algorithm MMseqs2. The performance of ESM-2 embeddings were not statistically significant compared to Foldseek both at top-1 rank and any rank comparisons, whereas both structure-homology derived and embedding-derived function transfer were substantially better than sequence-derived calls.

These examples illustrate the funnel tiers, and using the retrospective DUFs benchmark, we quantified how many graded hits are unrelated to the query’s functional annotation as identified by our pipeline, representing the overall transferable error rate for prospective use. Our partitioning strategy for confident hits from queries with qTM ≥ 0.5 into the three tiers mentioned above thus separates the yield loss from the accuracy loss. Of 264 confident retrospective DUF queries, 151 (57.2%) yielded informative or specific claims, while the known domain controls arm did better, with 294/440 confident queries yielding a correct call. Among the claims made, the errors are predominantly overgeneral (at the root of the GO term) rather than unrelated. Across all 148 claims for the retrospective DUFs, 28 were unrelated (18.9% error), and overgeneral claims contribute to 60.3% of the answers obtained.

The known domain controls arm behaves the same way: of the 294 confident queries, 66% of the claims are overgeneral, and 15% are returned as unrelated (**Supplementary Table S3**). If we split the data and looked only at ‘confident + informative’ and ‘confident + specific’ strata, the unrelated rate for retrospective DUFs was 8.6% and 21.2% (error rate), respectively, whereas for the known domain controls arm, the unrelated claims were 3.6% and 16.2%, respectively (**Fig. 3B**). Comparing the retrospective DUFs and the known domain control arms, the unrelated rate for the DUF arm is 1.3 times that of the control, which is expected if the DUF arm’s difficulty lies in the specificity of the available annotation rather than the structural search. Alongside Foldseek searches for structural homology, we also evaluated the same benchmark queries from both the retrospective DUFs and the known domain control arms using a strictly sequence-based homology search (MMseqs2) and the ESM-2 protein language model to corroborate Foldseek findings.

Against MMseqs2 run on the same 648 attempted queries per arm, Foldseek retrieved a surviving hit for 566 retrospective DUF queries *vs*. 217 for MMseqs2, which is a ∼2.6-fold difference in retrieval rate, and made a top-1 call for 13.9% of queries versus 5.1%. On the known domain control arm, Foldseek returned hits for 599 queries versus 314 for MMseqs2, with a top-1 call of 27.8% and 17.7% for Foldseek and MMseqs2, respectively, showing Foldseek consistently outperforming MMseqs2 on both arms (McNemar exact test, retrospective DUFs top-1 *P* = 1.3 x 10^-12^, known domain controls *P* = 1.2 x 10^-7^). On examining hits for which both Foldseek and MMseqs2 had a rank-1 call, the precision is 15.9% for Foldseek and 15.2% for MMseqs2 for the DUF arm, whereas 30.1% *vs*. 36.6% for the known domain control arm. When a top-1 rank is available, the sequence method is nominally more precise; however, structural searches win by finding something to transfer more often than the sequence-only searches (**Fig. S3D**) (**Supplementary Table S4**).

In addition to Foldseek and MMseqs2 searches on the benchmark dataset, we also incorporated ESM-2 protein language model to search a subset of 94 DUF queries drawn from families filtered to avoid ESM training contamination and restricted to those with usable ground truth (**Supplementary Table S5**). Comparing Foldseek and ESM-2 embedding search side-by-side, Foldseek’s top-1 correct rate was 24.5% (23/94) against 18.1% (17/94) for an embedding search, and the two were indistinguishable at any rank (37.2% each, 35/94) (McNemar exact test, top-1 rank *P* = 0.33; any-rank *P* = 1). Matching candidate depth at 100 candidates (to compare at equal output depth with ESM-2 calls), Foldseek’s top-1 rate dropped to 20.2%, showing both approaches are complementary and their union reaching 52.1% (49/94) informative calls from the searches (**Fig. 3C**). Overall, comparing three different approaches informed Foldseek as a reliable and easy-to-use method for large-scale structural homology searches.

### Calibrated structural homology transfer yields putative molecular functions for still DUFs

After calibrating structural homology transfer on DUF families whose functions were later established, we applied it prospectively to Pfam DUF families that remain functionally uncharacterized (still DUFs). Of the 344 still DUF families (**Fig. 1C**) in four model proteomes, 296 query domains were extracted and folded across the four proteomes, and 262 returned at least one structural hit after family- and clan-level masking, covering 218 distinct Pfam families. At the calibrated cut-off of query-anchored TM-score ≥ 0.6, 98/262 query domains (37.4%) returned a confident structural match in at least one database (**Fig. 4A**). Of these, 35 (13.4% of 262 queries) matched a target carrying an informative molecular function annotation, and 27 (10.3%) matched a target whose annotation was specific rather than a general parent term.

**Figure 4:**
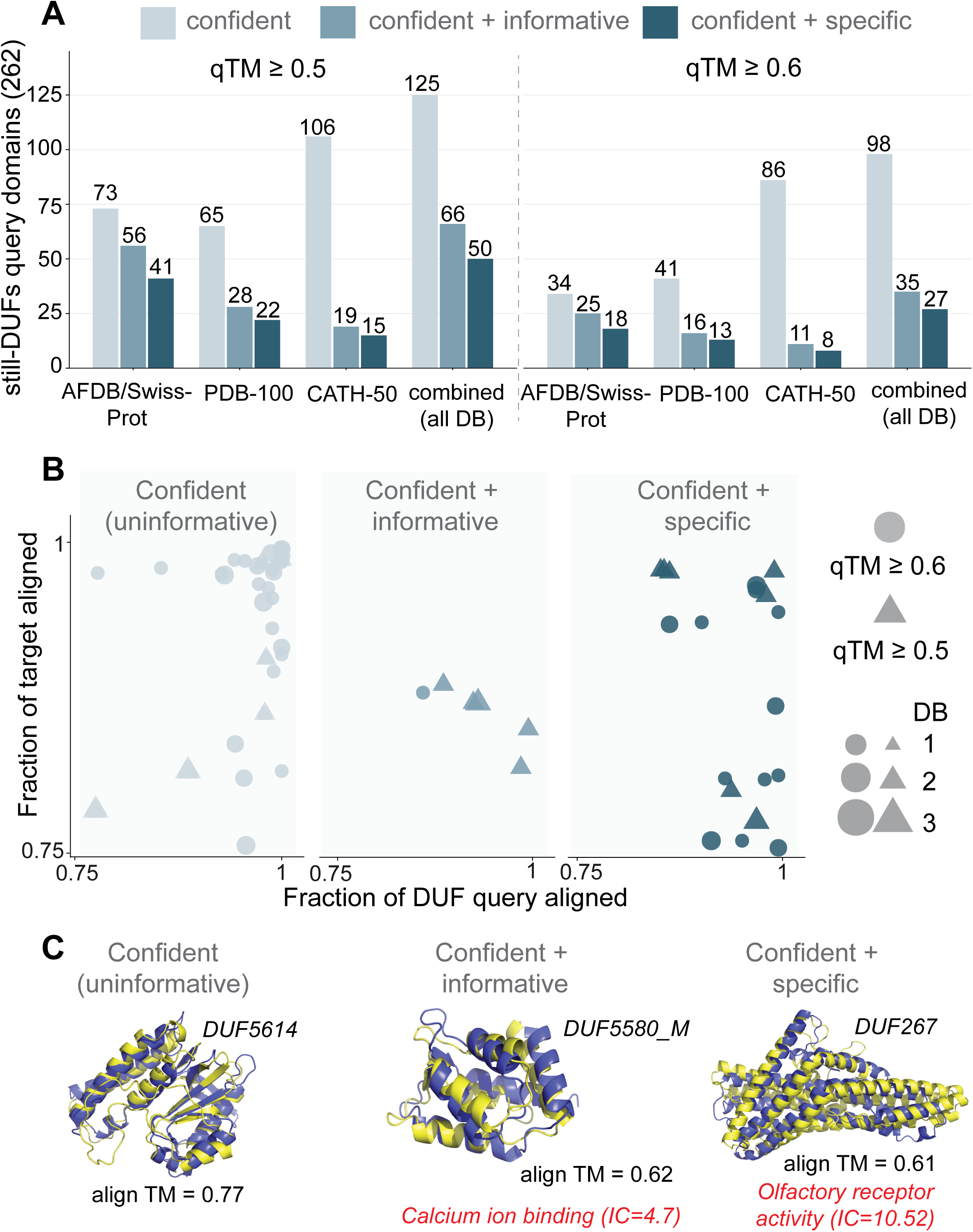
Structural homology searches on still DUFs and classification of hits to three funnel tiers. A) Distribution of Foldseek-derived hits to three funnel tiers (confident, confident+ informative, and confident + specific) for 262 evaluable still DUFs against three databases (AFDB/SwissProt, PDB-100, and CATH50), separated into two different cut-offs at qTM 0.5 and 0.6. B) Fraction of query and target alignment for three funnel tiers with support from union of hits from multiple databases, separated by two different qTM cut-off values. C) Representative examples of hits from three funnel tiers, where confident only hits are uninformative with no molecular function transfer, whereas the confident + informative and confident + specific tiers are separated by the information content (IC) scores, with scores < 7 tiered to confident + informative, while IC ≥ 7 were grouped in the confident + specific class.

Relaxing the cut-off of query-anchored TM-score ≥ 0.5 raised these to 125 confident (47.7%), 66 informative (25.2%), and 50 specific (19.1%), when all three databases were combined (**Fig. 4A**). Our structural homology searches find that the databases were markedly non-interchangeable. CATH-50 produced by far the most confident matches (86/98 at qTM ≥ 0.6, 46 of them found in no other database) but the fewest specific calls (8, only 2 uniquely; **Supplementary Table S6**). On the other hand, AFDB/SwissProt produced fewer confident matches (34, only 2 uniquely) but the most specific calls (18, 11 uniquely), while PDB100’s performance was intermediate (41 confident, 8 unique; 13 specific, 5 unique). Combining three different search databases revealed that a domain-level fold assignment and a transferrable function are two different retrieval problems: the database that best recognizes the fold is the one least able to name what it does (as a specific function), and searching all three and taking the union recovered 27 specific calls where the best single database (AFDB/SwissProt for functional annotation transfer) recovered 18 (**Fig. 4A**).

When the TM-score cut-off was relaxed, at qTM ≥ 0.5, 59/125 confident query domains matched targets, and these were not marginal matches. The best hits had a higher median qTM-score than the domains that did yield a call (0.725 *vs*. 0.606, Mann-Whitney *P* < 0.0001) (**Fig. S4A**), with higher sequence identity (median fident 0.230 *vs.* 0.133) and near-complete coverage of the target as well as the query (median target coverage 0.891 *vs.* 0.460). In other words, despite high query and target alignment score of > 0.75 for several proteins supported by hits from multiple databases (**Fig. 4B**), the structurally cleanest still DUF matches were disproportionately to other uncharacterized proteins. Indeed, our searches against the AFDB/UniProt50 database comparison also revealed that the failure to obtain molecular function annotation is due to target annotation state rather than from masking (**Fig. S4B**). Nevertheless, relaxing the TM-score cut-off from 0.6 to 0.5 adds 23 specific calls, which is a substantial fraction of the yield. So, we asked the benchmark calibration arms how much relaxing the TM-score would cost. In the known domain controls arm, the error rate—which are the confident calls naming a function unrelated to the true one rose from 13.1% (29/221) when qTM ≥ 0.6 to 31.1% (14/45) with a relaxed qTM ≥ 0.5. Therefore, while we report still DUF results at both qTM ≥ 0.5 and qTM ≥ 0.6 cut-offs, the results obtained from a stringent cut-off are more reliable based on the benchmark error rates. In **Fig. 4C**, we provide three illustrative examples of structural homology for each of the three categories (confident, confident + informative, and confident + specific) based on the IC score for the still DUFs.

For the informative calls (confident + informative; **Fig. 4A**) at qTM ≥ 0.6 (n=35), we asked whether other orthogonal evidence for function may be obtained from UniProt, Prosite, or PANTHER orthology. None of the 35 calls were supported by a UniProt ACT_SITE or BINDING annotation. On the other hand, Prosite and SMART/CDD together identified sequence profiles for 8/35 calls, whereas PANTHER orthology assigned 22/35 to a family, though often naming a paralog group than a molecular function. Taken together, 22 of 35 ‘confident + informative’ calls had orthogonal sequence-derived support. However, despite orthogonal evidence for putative function, we argue that since most of these calls rest on structural evidence, they require experimental follow-up rather than additional computational confirmation. In addition to orthogonal methods to infer putative molecular functions, we also used the Mechanism and Catalytic Site Atlas (M-CSA) (45) database to examine whether catalytic residues are conserved for calls that fall within an annotated enzyme class.

Because alignment score alone discards genuine homologies below a cutoff (qTM ≥ 0.5 or 0.6), we examined whether conservation of annotated catalytic residues inside the aligned space carried additional information even when hit with qTM < 0.3 was considered. For the benchmark query sets (retrospective DUFs and known domain controls), we find that among control hits with qTM < 0.3, calls in which at least one annotated catalytic residue was conserved were correct 57.9% of the time (11/19) against 17.3% (9/52) when the site was present but not conserved (Fisher’s exact test *P* = 0.002). While the same contrast held at qTM ≥ 0.5 (92.3%, 36/39 vs 41.2%, 7/17; Fisher’s exact test *P* = 0.0001) for the known domain controls, in the retrospective DUF arm, the enrichment was not significant when qTM < 0.3 hits were considered or with higher TM-scores of qTM ≥ 0.5 (**Fig. S5**). We therefore treat catalytic-residue conservation as a promising but unvalidated filter for still DUF candidates. Although the known domain control arm establishes the principle, the DUF arm counts are too small and too enzyme-poor to support robust, specific calls for candidate enzyme hits (**Supplementary Table S7**).

### A candidate prioritized set of still DUFs to facilitate experimental studies

Applying both relaxed and strict qTM cut-offs of 0.5 and 0.6, along with multi-database corroboration, yielded a shortlist of 50 still DUF domains with a ‘confident + specific’ call to facilitate experimental follow-up studies (**Supplementary Table S8**). This includes both conserved as well as lineage-specific DUF families. **Fig. 5** illustrates examples of lineage-restricted DUF families, such as DUF7778 (*C. elegans*), DUF4777 (*D. melanogaster*), and DUF 5580_C (*M. musculus*). DUF7778 represents a pleckstrin-homology (PH domain fold with a GO:0141038 functional prediction of kinase activator activity (**Fig. 5A**). PH domain is an ∼100 amino acid domain found in several human proteins involved in cell signaling and as constituents of the cytoskeleton (46), and consists of two perpendicular anti-parallel beta sheets, followed by a amphipathic helix (47). While PH domains occur in a variety of contexts, it is widely recognized to facilitate protein-protein interactions (48). Similarly, DUF4777—a lineage-restricted DUF found predominantly in insects represents a winged-helix fold, with an assigned GO: 0003691 associated with DNA binding activity (**Fig. 5B**). Winged-helix domain (WHD) is widespread across all domains of life with its known nucleic-acid binding roles (49). WHD has a characteristic domain architecture composed of a helix-turn-helix motif (HTH), which has two helices connected by a sharp turn, and are widespread double-stranded DNA binding element found in ∼30% of DNA-binding proteins (50). Despite a molecular function transfer of DNA binding activity, it is also worth noting that WHDs also mediate protein-protein interactions (51).

**Figure 5:**
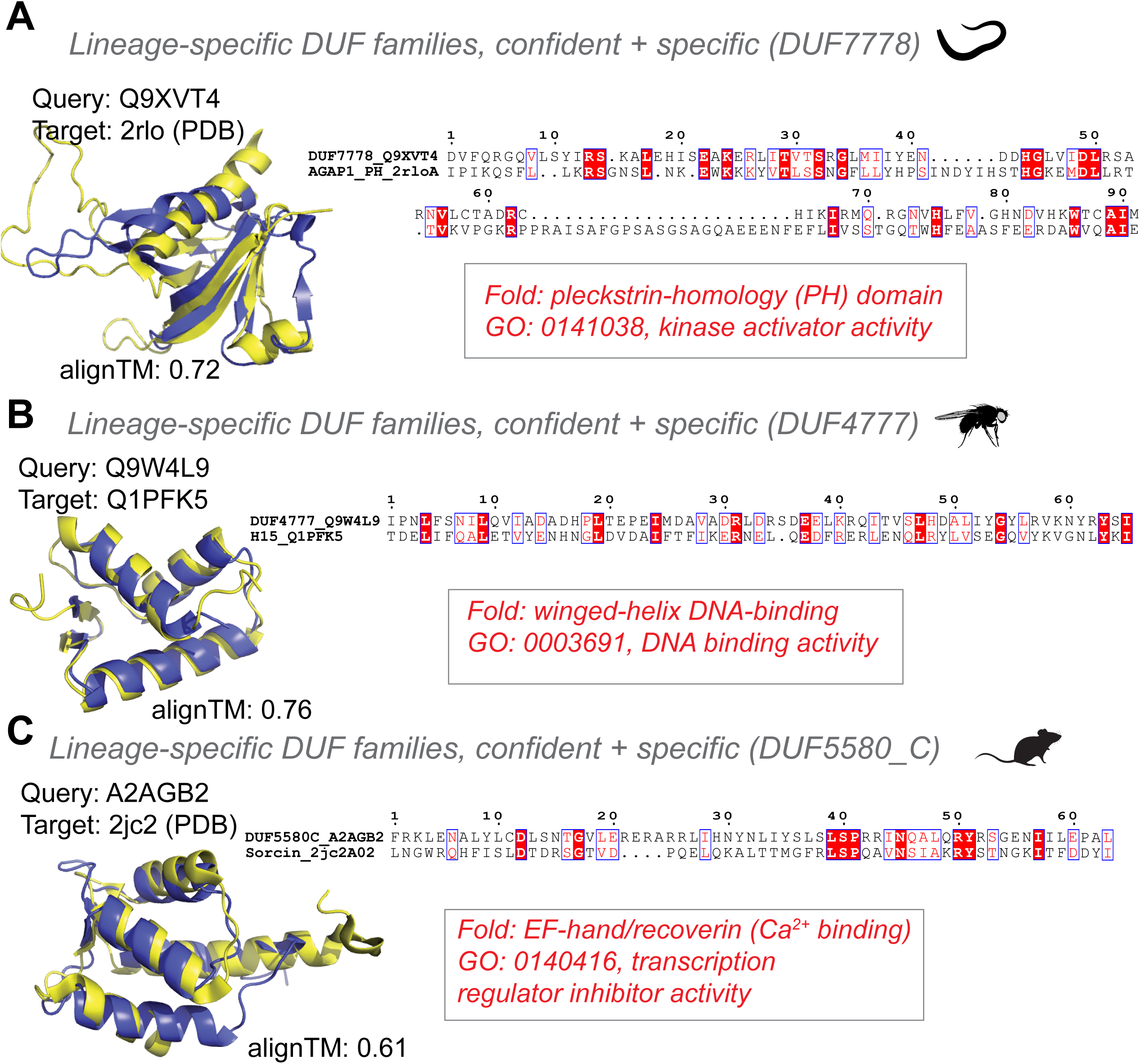
Examples of lineage-specific DUF families with confident molecular function transfer. Three illustrative examples of confident + specific DUF family molecular function transfer are shown for A) DUF7778 in *C. elegans* corresponds to a pleckstrin-homology (PH) domain, with a GO transfer (GO: 0141038) of predicted kinase activator activity. B) DUF4777 is an insect specific DUF family and the structural homology search identified a winged-helix domain fold for this family with a GO molecular function (GO: 0003691) of predicted DNA binding activity. C) Similarly, DUF5580_C in *M. musculus* recovered EF-hand/recoverin fold, with a predicted transcription regulator inhibitor activity (GO:0140416).

Finally, DUF5580_C is a EF-hand/Recoverin (Ca^2+^ binding) with a GO:0140416 term associated with transcription regulator inhibition activity (**Fig. 5C**). EF-hand Ca^2+^ binding proteins are involved in the modulation of Ca^2+^ signals and direct transduction of ionic signal to downstream biochemical processes (52). EF-hand motif is widespread in animals (53) and are typically organized in structural domains containing two or more EF hands that form highly stable helical bundles (54). In addition to the illustrated examples, other representative candidates include DUF2371, which matched to ATP synthase subunit C in all three databases, DUF1182, matching soluble cytochrome B562, and DUF4791, matching a Wntless-like transmembrane domain (**Supplementary Table S8**). A point worth noting is that sequence identity across these alignments was 9-24%, so none would have been recovered by a sequence-level search. Furthermore, the 46 families behind the ‘confident + specific’ calls were taxonomically heterogeneous, spanning 25 to 17,301 taxa (median 683). 26/46 were restricted to Metazoa, and only 7 had bacterial representatives, and many of these DUFs were enriched for lineage-restricted families.

## Discussion

Understanding DUFs and resolving their molecular functions has remained a topic of interest (55, 56) ever since they were first introduced to biological lexicon nearly 25 years ago (57). In this work, we employed structural homology searches to identify putative molecular functions for DUFs in four model proteomes. In doing so, we set out to quantify, rather than assume, how reliably structural homology transfers molecular function to DUFs. Using families that were DUFs in Pfam 28.0 and have since been named in the latest release as a time-split benchmark, we find that Foldseek-based transfer recovers the Pfam 30.0-assigned function for 26.1% of retrospective DUFs with a structural hit, compared with 35.2% for the difficulty-matched known-domain control arm. When considering hits with a qTM ≥ 0.5 that assert something informative, we find that 55.4% are incorrect. This suggests that structural searches find genuine, non-trivial functional relationships for a minority of DUFs; however, slightly more than half of the confident assignments are incorrect, driven by overgeneral assignments where the transferred term is in the right functional neighborhood but too broad to constitute an answer.

We also lay out the importance of circularity masking and methodological findings from that exercise. Without masking, top-1 agreement is 84.7% in the DUF arm and 84% in the control arm, which is an excellent result, with no difference between well-characterized known domains and DUFs. However, masking the query’s own Pfam family and its clan siblings removes 43% of hits in the DUF arm and 59% in the control arm. Thus, the same data supports either an apparent success rate of 85% or a real one of 26%, depending on a single preprocessing decision. We also note that masking penalizes the control arm more, as well-characterized families at the structural-level (58), when grouped by evolutionary relationships (59) have more annotated clan-level relatives available to retrieve. Therefore, an unmasked pipeline inflates the easier arm more, which, by circular construction, can lead to an incorrect conclusion that DUFs perform on par with known domains.

From our time-split benchmark, the key question is why the error rate is so high, given that the structural alignments themselves are good (qTM ≥ 0.5). The mechanism is not retrieval failure but transfer failure in obtaining molecular function. Errors are dominated by overgeneral assignments rather than unrelated ones (67% of errors are at the informative tier, but not specific). The overgeneral assignment is an expected consequence of fold-function decoupling, which is well documented for homologous enzyme superfamilies (35), and it also means that the error rate is sensitive to how much one demands of a claim. If we strictly considered only specific-tier assignments, the DUFs error rises to 60.2% and accepting general-tier findings lowers it to the reported 55.4%. It is important to note that general-tier terms carry a median information content (41) score of 5.34, compared with 10.8 for the specific-tier category, indicating that lower error comes at the cost of weaker claims for molecular function.

A related and initially counterintuitive result is that increasing the qTM cutoff to higher values did not buy accuracy. Increasing the qTM cut-off from 0.5 to 0.7 discards 83% of informative calls (66 to 11) while reducing the benchmark error rate only by 5% points, from 55.4% to 50.8%. The implication of this result is that more structural evidence does not make the functional inference more likely to be correct. In other words, structural confidence and annotation-transfer are different quantities and a threshold on the former is not a proxy for the latter. Nevertheless, compared to sequence search, structural homology searches deliver a large and unambiguous advantage, as demonstrated by a recent study in predicting the roles of DUFs involved in protein-protein interactions (60). In the retrospective DUF arm, Foldseek recovers the correct function for 13.9% of attempted queries against 5.1% for MMseqs2 and the advantage comes from more coverage from structural searches, where 566/648 DUF queries returned a hit versus 217 for sequence search.

Furthermore, with our limited set of evaluable queries, the comparison between protein language model embeddings and Foldseek was not distinguishably different. Although embeddings are generally faster, side-by-side evaluations like ours have concurred that embeddings cannot replace structural alignments, but can greatly help improve their quality (61, 62). On the 94 queries where both methods were evaluable, Foldseek achieved 24.5% top-1 accuracy against 18.1% for embedding-based transfer—a difference that is not significant (McNemar exact test *P* = 0.33), and the two methods were identical at 37.2% when any rank was considered (instead of just top-1). It is important to note that, with a small query set, we are underpowered for this comparison. Therefore, the unbiased evaluation is that we cannot separate the two, rather than claiming they are equivalent, and a future study that seeks to establish structural search as superior to embedding-based transfer would require a substantially larger paired sample than the ones we compared in this study.

Several other limitations are worth discussing here. First, our ground truth is Pfam curation, so we measure agreement with curated entries rather than known experimental answers. Families named entirely on computational evidence can therefore reinforce the same inferences made by our pipeline, potentially inflating its apparent performance. Second, the families named since Pfam 28.0 are unlikely to represent a random sample of DUFs; they were probably among the most tractable, which may inflate our estimated performance. Thus, the observed error rate should be regarded as a lower bound for families that remain unnamed. Third, and most importantly, our assignments lack experimental validation. An attempted orthogonal structural assessment provided limited support, and cross-database comparisons were possible for only 13 of 35 confident, informative assignments. Individual assignments should therefore be interpreted as hypotheses with an estimated error rate, not as validated findings. Fourth, for a substantial fraction of still DUFs, no reliable molecular function transfer could be obtained despite good structural alignment. To tackle this, recent methods that integrate structural and evolutionary approaches (63) for domain annotation may prove fruitful in refining the annotation of challenging DUFs. Finally, because our benchmark and query sets are limited to four model eukaryotes, it remains unclear whether these performance estimates generalize to prokaryotes or to the much larger set of metagenome-derived families, where structural coverage and annotation density differ substantially.

Overall, our results suggest a shift in how structural DUF assignments should be used. Applied to 296 unannotated families, our calibrated pipeline produced 50 confident, specific assignments, approximately half of which are expected to be correct. Although this confidence level is insufficient to support strong conclusions about any individual family, it provides a valuable framework for prioritizing experimental validation. A 45% probability of correctness represents a substantial improvement over randomly selecting an uncharacterized family, while the explicit error estimate allows researchers to plan validation efforts accordingly. Thus, the current value of structure-first annotation lies not in closing the annotation gap computationally, but in making that gap experimentally tractable by ranking candidates. The next step is to evaluate this calibration against experimental rather than curatorial outcomes, either by testing a subset of our candidates or by repeating the benchmark in using families subsequently characterized through experiments.

## Materials and Methods

### Time-split benchmark construction

The retrospective benchmark in our study comprises DUF families whose functions were unknown in Pfam release 28.0 but have since been resolved and named in Pfam 32.0 (n=353 across 4 focal species). Families absent from the current release (retired or merged, n=49) were dropped due to lack of ground truth for further assessments. We then intersected the resulting family list with the UniProt reference proteome (*S. cerevisiae*: UP000002311, taxon 559292; *C. elegans*: UP000001940, 6239; *D. melanogaster*: UP000000803, 7227; *M. musculus*: UP000000589, 10090) to confirm that every benchmark query is a protein annotated in the organism. For each retrospective DUF family, we identified one protein per family per organism using the following criteria: the protein’s reviewed status, the presence of an AlphaFold model, and then the UniProt annotation score. Similarly, we identified controls, that are Pfam families with a known function to complement the retrospective DUF set drawn from the same proteome and matched 1:1 to the DUF queries using domain length and the mean per-residue pLDDT as criteria. A list of UniProt IDs for retrospective DUFs and matched known domain controls are provided in **Supplementary Table 1**. For these queries, we fetched domain boundaries per protein from the InterPro API, and AlphaFold PDB structures and mmCIF files from the AlphaFold database.

### Structural homology search

Structural homology searches were run with *foldseek easy-earch* using a fixed output column set with the following parameters: *--alignment-type 2 -s 9.5 --max-seqs 2000 -e 10--threads 16 --exhaustive search*, with the following requested columns: *query, target, fident, alnlen, mismatch, gapopen, qstart, qend, tstart, tend, evalue, bits, prob, alntmscore, qtmscore, ttmscore, lddt, qlen, tlen, qaln, taln*. For target databases, we used Foldseek’s prebuilt databases, AFDB/SwissProt, PDB100, CATH50, and AFDB/UniProt50. Top 5 candidates were retained per query per database after circularity masking (see below) and the default candidate ranking is by e-value; however, we also conducted ranking by alignment TM-score, and was evaluated as a sensitivity analysis, and we report results from e-value ranking.

### Circularity control: family and clan-level masking

Besides self-hits, searches for DUF families typically return self-family (the same DUF) as the top hits. To remove circularity, we applied masking at both family and clan level. The clan-level hits are the structural answer under a different accession, and family-only masking leaves the circularity partly intact. The family- and clan-level information was obtained from Pfam, and masking was applied identically to the retrospective DUFs and the known domain controls. Note that masking removes a known domain control’s own correct answer as well, and we ensure that the retrieval for the positive control is verified before masking.

### Ground truth for the retrospective DUFs arm

We checked the ground truth match for the retrospective DUFs using GO molecular function terms for the query family, resolved in two tiers. The first tier used the InterPro2Go mapping (https://www.ebi.ac.uk/GOA/InterPro2GO), and the fallback for families with no InterPro2Go mapping used the GO-MF terms recovered from reviewed proteins that carry the family. Failure to obtain information from both tiers was recorded as none/no functional match. Agreement between a query family and the hit’s annotation was evaluated on the GO-directed acyclic graph and not by string matching. Both the query and the target sides were ancestor-closed and only restricted to the molecular function aspect. Information content (IC) was then estimated as -log2 of the frequency of terms, and IC was binned into three groups: Specific (IC ≥ 7), General (IC ≥ 3), and Wrong (IC below 3 or lacking usable GO-MF annotation). The IC scores for the three categories were calibrated against 1480 semantic adjudications across 562 pairs in which both sides carried a GO term. Mean shared-ancestor IC from this set for calibration was 2.0 (Wrong), 4.34 (General), and 5.83 (Specific). Although IC provides a stratification of hits, we recommend further manual inspection as a curation step to validate the results.

### Scoring for still-DUF category

For DUF families whose molecular function remains unknown, there is no ground truth, so agreement cannot be measured. In this case, we computed scores using what-hit asserts and classified hits that carry a molecular function term as specific if IC ≥ 7 and as general if IC ≥ 3 and < 7. For DUF families that registered hits with no GO molecular function, or carried only near-root terms were classified as uninformative. It is worth noting that the scoring property for the still-DUF category is a yield measure and not an accuracy measure.

### Catalytic residue transfer test

A high TM-score establishes shared shape but not retained chemistry, and the common failure mode of structural function transfer is a relative that shares the fold and loses the active site. For a set of still DUF family hits for which information is available on UniProt about active-site and binding features on the target, we examined residue conservation. We complemented this analysis with M-CSA (Mechanism and Catalytic Site Atlas) data to examine conserved residues. Although M-CSA data is limited in scope, as only 988 reference proteins for which active site information was available in our case (0.39% of 11,201 target accessions), data from M-CSA added informative conservation calls for 126 positions across 44 M-CSA proteins that occur in our targets. To conduct M-CSA conservation analysis, we mapped the target residue numbers to the query, and conservation requires both alignment and amino acid identity. Each hit was graded as i) conserved, ii) partial, iii) lost, and iv) unalignable.

### Additional database and baseline arms

For the DUF queries, we also used sequence-level search using MMseqs2. Target sequences were extracted from the same foldseek database via foldseek *convert2fasta* (590,183 entries, AFDB/SwissProt). Query sequences are the same per-domain PDB files foldseek searched, and the query lengths were verified for all 1296 queries. MMseqs2 was run using the following command: *mmseqs easy-search, -s 7.5, -e 10.0, --max-seqs 2000, --num-iterations 3*. In addition to MMseq2, we also searched the same target universe with ESM-2 embeddings rather than structural or sequence alignment. Because ESM-2 was trained on UniRef50 with a September 2021 cutoff, queries were first filtered for training contamination: of the 354 DUF families in the benchmark,109 that were still explicitly DUF-named in Pfam 35.0 (November 2021, i.e. after the model cutoff) were retained, yielding 174 query domains of which 94 have usable ground truth. Queries and targets were embedded with ESM-2 650M (*facebook/esm2_t33_650M_UR50D*). Target sequences (534,236 proteins, 191.8 M residues) were extracted from the same Foldseek database the structural arm searched, so that the three arms searched identical target sets rather than merely similar ones. Each protein was represented by the mean of its final-layer residue representations. Retrieval took the top 100 targets per query by cosine similarity, computed in chunks of 20,000 targets with a partial sort. The resulting hits were scored through the main pipeline’s annotation, self-hit masking and GO-scoring functions.

### Pipeline execution and availability

The pipeline is provided as a self-contained Python package that can be executed on the command-line, which requires: pandas == 3.0.5, numpy == 2.5.1, and matplotlib. The ESM arm requires pytorch and transformers. Along with that, we also provide a user-friendly python notebook to conduct searches on user-defined DUF queries against AFDB/SwissProt, and AFDB50, where the Jupyter book pipeline ranks hits using lexical terms associated with GO molecular function. The pipeline and associated scripts are available for download at Zenodo: DOI: 10.5281/zenodo.22693246.

